# Viral mediated tethering to SEL1L facilitates ER-associated degradation of IRE1

**DOI:** 10.1101/2020.10.07.330779

**Authors:** Florian Hinte, Jendrik Müller, Wolfram Brune

**Affiliations:** Heinrich Pette Institute, Leibniz Institute for Experimental Virology, Hamburg, Germany

## Abstract

The unfolded protein response (UPR) and endoplasmic reticulum (ER)-associated degradation (ERAD) are two essential components of the quality control system for proteins in the secretory pathway. When unfolded proteins accumulate in the ER, UPR sensors such as IRE1 induce the expression of ERAD genes, thereby increasing protein export from the ER to the cytosol and subsequent degradation by the proteasome. Conversely, IRE1 itself is an ERAD substrate, indicating that the UPR and ERAD regulate each other. Viruses are intracellular parasites that exploit the host cell for their own benefit. Cytomegaloviruses selectively modulate the UPR to take advantage of beneficial and inhibit detrimental effects on viral replication. We have previously shown that murine and human cytomegaloviruses express homologous proteins (M50 and UL50, respectively) that dampen the UPR at late times post infection by inducing IRE1 degradation. However, the degradation mechanism has remained uncertain. Here we show that the cytomegalovirus M50 protein mediates IRE1 degradation by the proteasome. M50-dependent IRE1 degradation can be blocked by pharmacological inhibition of p97/VCP or by genetic ablation of SEL1L, both of which are component of the ERAD machinery. SEL1L acts as a cofactor of the E3 ubiquitin ligase HRD1, while p97/VCP is responsible for the extraction of ubiquitylated proteins from the ER to the cytosol. We further show that M50 facilitates the IRE1-SEL1L interaction by binding to both, IRE1 and SEL1L. These results indicate that the viral M50 protein dampens the UPR by tethering IRE1 to SEL1L, thereby promoting its degradation by the ERAD machinery.

**Importance:** Viruses infect cells of their host and force them to produce virus progeny. This can impose stress on the host cell and activate counter-regulatory mechanisms. Protein overload in the endoplasmic reticulum (ER) leads to ER stress and triggers the unfolded protein response, which in turn upregulates protein folding and increases the degradation of proteins in the ER. Previous work has shown that cytomegaloviruses interfere with the unfolded protein response by degrading the sensor molecule IRE1. Herein we demonstrate how the viral M50 protein exploits the ER-associated degradation machinery to dispose of IRE1. Degradation of IRE1 curbs the unfolded protein response and helps the virus to increase the synthesis of its own proteins.

## Introduction

The ER is a cellular compartment responsible for synthesis, assembly, and trafficking of secretory and membrane proteins (1). To maintain protein homeostasis, cells must ensure proper protein folding and maturation. When the load of newly synthesized proteins and the folding capacity get out of balance, unfolded and misfolded proteins accumulate in the ER, resulting in ER stress (2). To maintain ER homeostasis, cells have evolved quality control systems and counter-regulatory mechanisms.

While protein aggregates in the ER are removed by autophagy, the primary mechanism for the disposal of ER-resident proteins is ER-associated degradation (ERAD) (3, 4). ERAD involves ubiquitylation and retrotranslocation of proteins from the ER to the cytosol, where proteasomal degradation takes place. The best-characterized and most conserved ERAD machinery in mammalian cells consists of the SEL1L-HRD1 protein complex and the transitional ER ATPase p97, also known as valosin-containing protein (VCP). Ubiquitylation is mediated by the E3 ubiquitin ligase HRD1 (also known as Synoviolin, SYVN1), which resides in the ER membrane and uses SEL1L as a cofactor (5). Ubiquitylated ERAD substrates are extracted from the ER membrane and translocated to the cytosol in an energy-dependent process mediated by the p97/VCP, a protein of the AAA (ATPases associated with diverse cellular activities) family (6).

Accumulation of unfolded or misfolded proteins in the ER triggers the unfolded protein response (UPR). It relies on three sensors, IRE1, PERK, and ATF6, which are activated upon ER stress and mediate signal transduction from the ER to the cytosol and the nucleus (reviewed in (7)). The PKR-like ER kinase (PERK) phosphorylates the translation initiation factor eIF2α, thereby reducing protein translation. The other two sensors, the inositol-requiring enzyme 1 (IRE1, also known as IRE1α or ER-to-nucleus signaling 1, ERN1) and the activating transcription factor 6 (ATF6), activate the transcription factors X-box binding protein 1-spliced (XBP1s) and ATF6(N), respectively. XBP1s and ATF6(N) stimulate the expression of chaperones, foldases, and components of the ERAD machinery. Thus, the three UPR sensors restore ER homeostasis by reducing the protein load and by increasing the protein folding capacity (7). Intriguingly, the UPR and ERAD regulate each other: the UPR sensors that upregulate ERAD (i.e., IRE1 and ATF6) are themselves subject to ERAD (8). IRE1 is recognized by the substrate recognition factor OS9 and is degraded by the SEL1L-HRD1 ERAD complex as part of its natural turnover (9). A similar mechanism has been described for ATF6 (10).

Viral replication within the host cell requires the synthesis of substantial amounts of viral proteins. Especially during the late phase of the viral life cycle, large quantities viral envelope glycoproteins and immunomodulatory transmembrane proteins have to be synthesized. This can overwhelm the folding and processing capacity of the ER and cause ER stress (11). Many viruses, particularly those of the *Herpesviridae* family, have evolved means to modulate the UPR and to exploit ERAD to their own benefit (reviewed in (12)).

Human cytomegaloviruses (HCMV, human herpesvirus 5) is an opportunistic pathogen and a leading cause of morbidity and mortality in immunocompromised patients. It is also the leading cause of congenital infections, which can result in long-term neurological deficits (13, 14). Murine cytomegalovirus (MCMV) is a related herpesvirus of mice, which serves as a small animal model for HCMV (15). During co-evolution with their respective hosts, the CMVs have acquired the ability to activate and regulate the UPR, as shown in pioneering work by the laboratory of James Alwine (16–19). Subsequent work by several laboratories has shown that HCMV and MCMV manipulate all three branches of the UPR (reviewed in (12)). For instance, we have recently shown that MCMV briefly activates the IRE1-XBP1 signaling pathway during the first few hours of infection to relieve repression by XBP1u, the product of the unspliced *Xbp1* mRNA. XBP1u inhibits viral gene expression and replication by blocking the activation of the viral major immediate-early promoter by XBP1s and ATF6(N) (20). At late times post infection, MCMV inhibits IRE1-dependent signaling by downregulating IRE1 levels to prevent deleterious effects of the UPR on virus progeny production. The MCMV protein M50 interacts with IRE1 and causes its degradation, a function it shares with its homolog in HCMV, UL50 (21). However, the precise mechanism of IRE1 degradation has remained unknown.

Herein we show that M50 reduces IRE1 protein levels by inducing its degradation via the proteasome. M50-mediated IRE1 degradation depends on p97/VCP and SEL1L, two component of the ERAD machinery. We further show that M50 interacts with both, IRE1 and SEL1L, suggesting that it functions as a viral adaptor which facilitates the interaction of the two proteins. These findings indicate that the viral M50 protein inhibits the IRE1 branch of the UPR by tethering IRE1 to SEL1L, thereby promoting its degradation by the ERAD machinery.

## Results

### M50 induces proteasomal degradation of IRE1

We have previously shown that M50 reduces IRE1 protein levels by reducing its stability (21). However, the mechanism of IRE1 degradation has not been resolved. Attempts to inhibit IRE1 degradation with lysosomal inhibitors were unsuccessful, suggesting that degradation does not occur by autophagy (21). However, since it has been shown that IRE1 is subject to ERAD during its natural turnover (9), we tested whether M50-induced IRE1 degradation can be inhibited with a proteasome inhibitor. Mouse embryonic fibroblasts (MEFs) were transfected with plasmids encoding Myc-tagged IRE1 and either FLAG-tagged full-length M50 or a C-terminally truncated M50 (1-276) mutant, which does not induce IRE1 degradation (21). Cell lysates were harvested 48 hours after transfection and analyzed by immunoblot. The proteasome inhibitor MG-132 or DMSO (vector control) was added during the last 5½ hours before harvesting. As shown in Fig. 1A, MG-132 treatment inhibited M50- induced IRE1 degradation.

**Figure 1.**
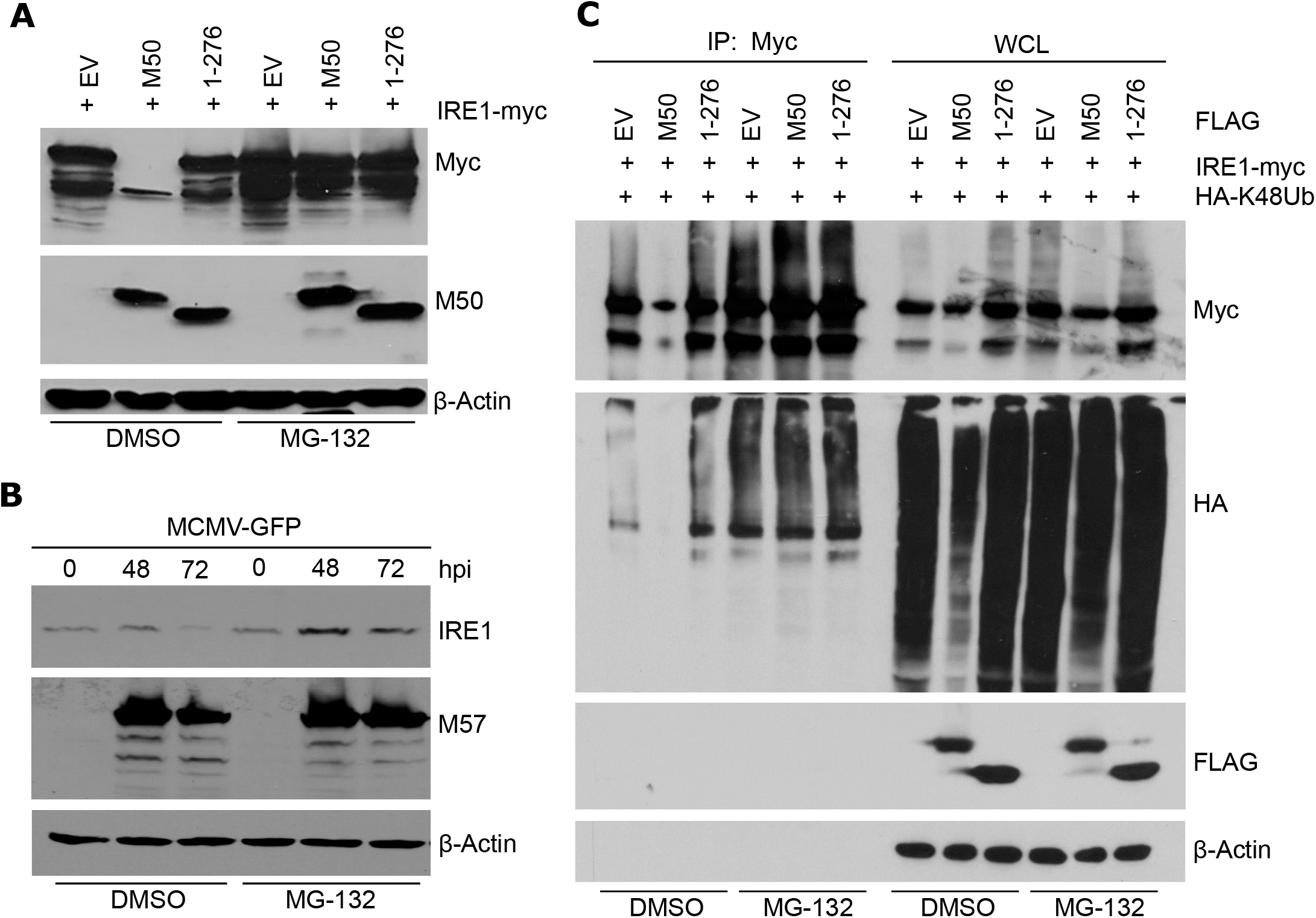
M50-mediated IRE1 degradation is blocked by proteasome inhibitor MG132. (A) MEFs were transfected with plasmids expressing myc-tagged IRE1 and full-length or truncated (1-276) M50 or empty vector (EV). Cell were treated with 25 μM MG-132 or DMSO for the last 5.5 hrs before harvesting of lysates at 48 hrs post transfection. (B) MEFs were infected with MCMV-GFP (MOI=3). Cells were treated with 30 μM MG-132 or DMSO for the last 6 hrs before harvesting. Cell lysates were analyzed by immunoblot. M57 was detected as an infection control. (C) MEFs were transfected with a plasmids expressing HA-tagged K48-only ubiquitin and plasmids as in A. Cell were treated with 30 μM MG-132 or DMSO for the last 6 hrs before harvesting of lysates at 48 hrs post transfection. IRE1 was immunoprecipitated with an anti-myc antibody. Ubiquitylated proteins in whole cell lysates (WCL) and in the immunoprecititated samples were detected by immunoblot with an anti-HA antibody.

Next, we tested whether MG-132 also inhibits IRE1 degradation in MCMV-infected cells, which occurs at late times post infection. MEFs were infected with MCMV-GFP, and IRE1 levels were analyzed at late times post infection. Treatment with MG-132 strongly increased IRE1 levels in MCMV-infected cells (Fig. 1B), suggesting that IRE1 degradation in infected cells occurred via the proteasome.

To test whether M50 induces proteasomal degradation of polyubiquitinated IRE1, we transfected MEFs with IRE1 and M50 expression plasmids and a plasmid expressing HA-tagged ubiquitin. A modified “K48-only” ubiquitin, which can only form K48-linked ubiquitin chains (22), was used in order to analyze the type of polyubiquitylation associated with proteasomal degradation. Myc-tagged IRE1 was immunoprecipitated, and ubiquitylated IRE1 was detected by immunoblotting. As expected, IRE1 overexpression resulted in a substantial level of polyubiquitylated IRE1 (Fig. 1C). In the presence of full-length M50, the level of polyubiquitylated IRE1 was massively decreased, and this effect was largely reversed when cells were treated with the proteasome inhibitor MG-132 (Fig. 1C). These findings suggested that M50 induces ubiquitylation and proteasomal degradation of IRE1.

ER-associated degradation of an ER membrane protein requires substrate recognition, polyubiquitination by an E3 ubiquitin ligase, and extraction from the ER membrane into the cytosol, where proteasomal degradation takes place. Extraction is mediated by the ATPase p97/VCP. Therefore, we used a specific p97 inhibitor to test whether M50-induced IRE1 degradation occurred via ERAD. In experiments analogous to those shown in Fig. 1A and B, the p97 inhibitor CB-5083 inhibited IRE1 degradation in M50-transfected as well as MCMV-infected cells (Fig. 2A and B).

**Figure 2.**
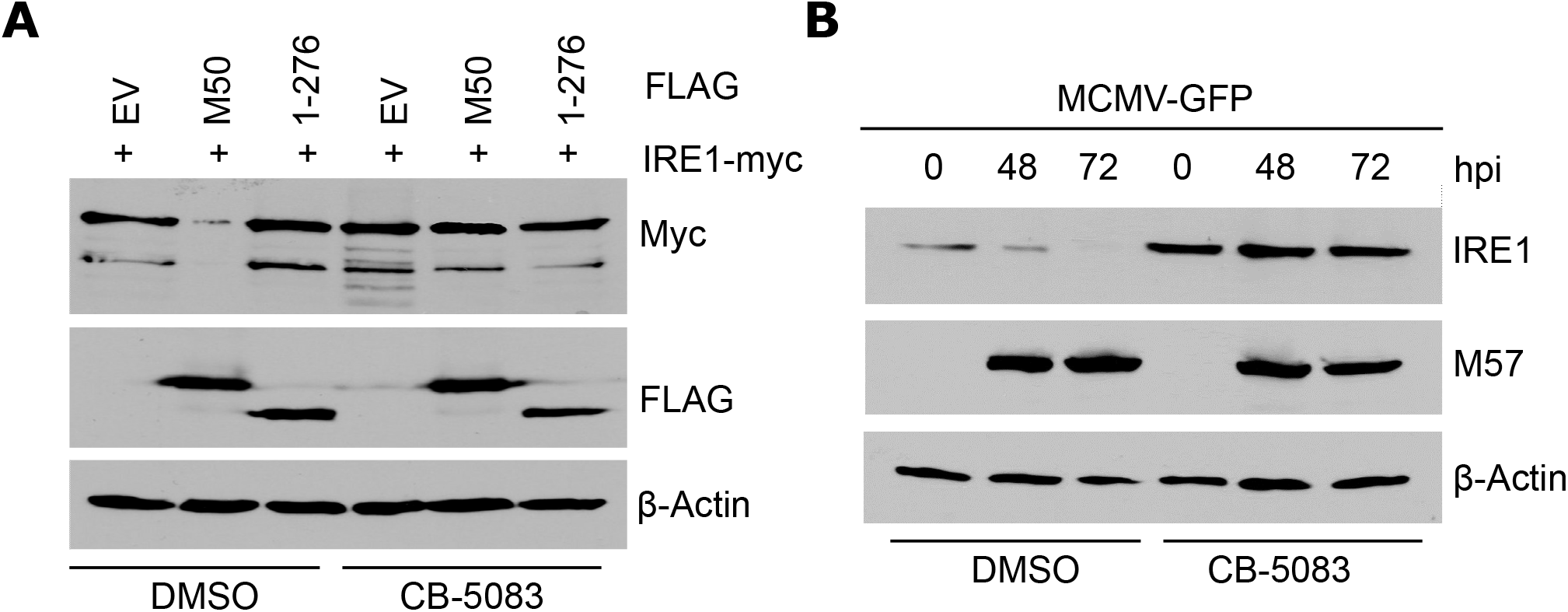
M50-mediated IRE1 degradation is blocked by p97/VCP inhibitor CB-5083. (A) MEFs were transfected with plasmids expressing myc-tagged IRE1 and FLAG-tagged full-length or truncated (1-276) M50 or empty vector (EV). Cell were treated with 30 μM CB-5083 or DMSO for the last 6 hrs before harvesting of lysates at 48 hrs post transfection. (B) MEFs were infected with MCMV-GFP (MOI=3). Cells were treated with 30 μM CB-5083 or DMSO for the last 6 hrs before harvesting. Cell lysates were analyzed by immunoblot. M57 was detected as an infection control.

### M50 interacts with SEL1L

During its natural turnover, IRE1 is recognized by the substrate recognition factor OS9, which functions as an adaptor protein of the SEL1L-HRD1 complex (9). We hypothesized that M50 could act as a viral adaptor protein that interacts with both, IRE1 and SEL1L. To test this hypothesis, HEK 293A cells were transfected with plasmids expressing Myc-tagged IRE1, FLAG-tagged M50, and untagged SEL1L. While IRE1 interacted only weakly with SEL1L in immunoprecipitation experiments, M50 interacted with both SEL1L and IRE1 (Fig. 3A). Expression of full-length M50 massively increased the interaction of IRE1 with SEL1L, whereas expression of the truncated M50(1-276) mutant did not (Fig. 3B). These results suggested that M50 acts as an adaptor protein that tethers IRE1 to SEL1L.

**Figure 3.**
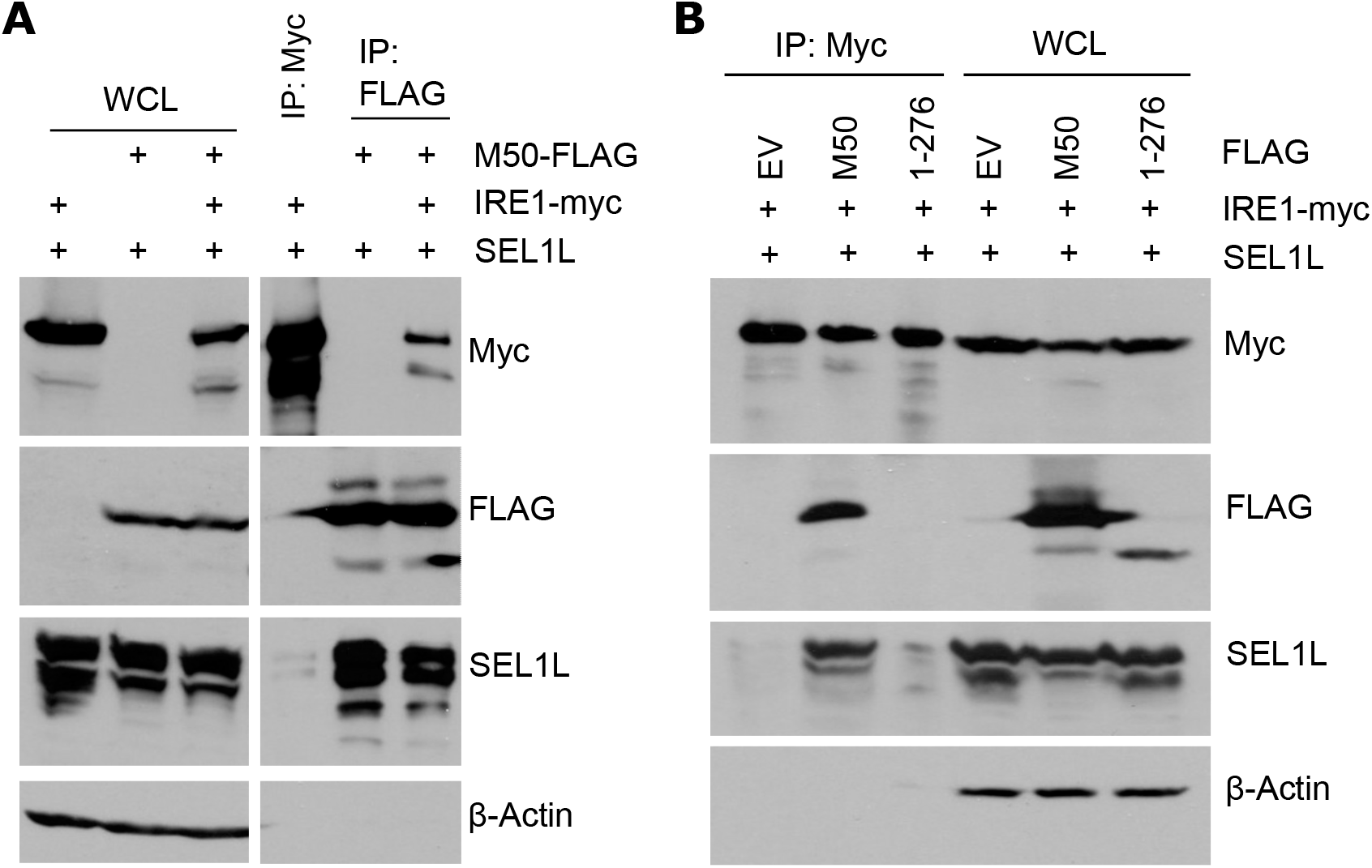
M50 interacts with IRE1 and SEL1L. (A) HEK 293A cells were transfected with a plasmids expressing SEL1L, Myc-tagged IRE1, and FLAG-tagged M50. IRE1-myc or M50- FLAG was immunoprecipitated. (B) HEK 293A cells were transfected with a plasmids expressing SEL1L, Myc-tagged IRE1, and FLAG-tagged full-length or truncated (1-276) M50 or empty vector (EV). IRE1 was immunoprecipitated with an anti-myc antibody. Proteins in whole cell lysates (WCL) and the immunoprecititated samples were detected by immunoblot.

### Genetic ablation of *Sel1L* abolishes M50-mediated IRE1 degradation

To test whether SEL1L is required for the M50-mediated IRE1 degradation, we generated *Sel1L* knockout MEFs by CRISPR/Cas9 gene editing. Two independent SEL1L-deficient cell clones were obtained with different guide RNAs (Fig. 4A). As SEL1L is involved in IRE1’s natural turnover, IRE1 levels are substantially increased SEL1L-deficient cells (9). Consistent with this published observation, we detected increased IRE1 protein levels in our *Sel1L* knockout MEFs (Fig. 4A). However, *Ire1* transcript levels were not significantly altered in these cells (Fig. 4B), confirming that impaired degradation rather than increased synthesis is responsible. Increased IRE1 protein levels correlated with increased *Xbp1* mRNA splicing and increased transcription of *Chop*, an XBP1s target gene (Fig. 4B). These results indicated that IRE1 protein levels and IRE1-mediated signaling are elevated in the absence of SEL1L.

**Figure 4.**
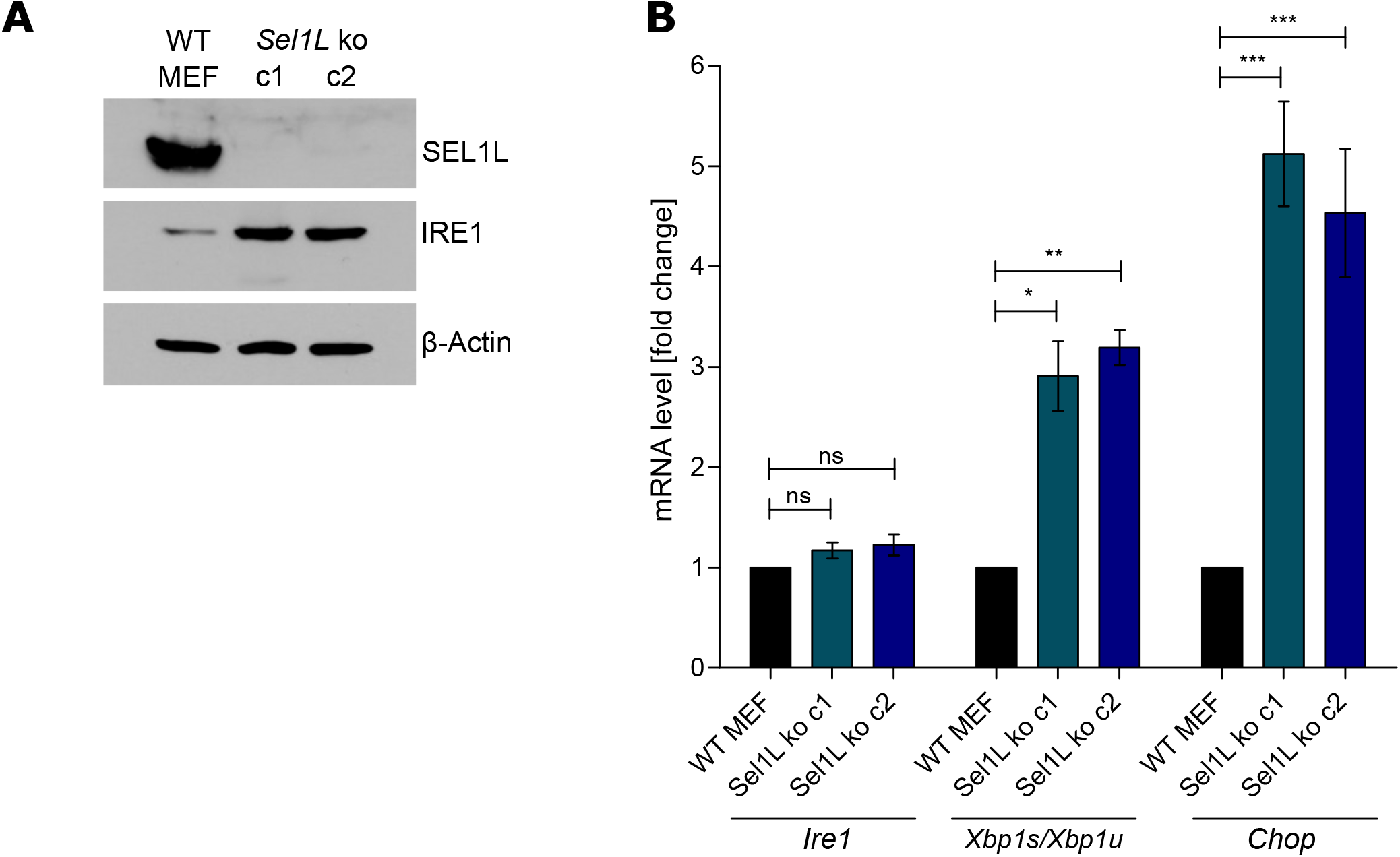
Generation of SEL1L-deficient MEFs. (A) *Sel1L* knockout MEFs were generated by CRISPR/Cas9 gene editing. Two independent clones obtained with different gRNAs are shown. SEL1L and IRE1 protein levels were analyzed by immunoblot. (B) Total RNA was isolated from WT and *Sel1L* knockout MEFs. mRNA levels of *Ire1, Chop, Xbpls*, and *Xbplu* were determined by qRT-PCR. *Gapdh* was used for normalization. Fold changes *(Sel1L* knockout compared to WT MEFs) are shown as means ±SEM of three biological replicates. *, p<0.05;***, p<0.01; ***, p<0.001.

Next, we tested whether the absence of SEL1L affects M50-induced IRE1 degradation. WT and *Sel1L* ko MEF were transfected with IRE1 and M50 expression plasmids. Expression of full-length M50 reduced IRE1 levels in WT MEF but not in SEL1L-deficient MEFs (Fig. 5A). Similarly, MCMV infection resulted in a strong reduction of IRE1 levels at late times post infection in WT but not in *Sel1L* ko MEF (Fig. 5B). We also analyzed IRE1 signaling in MCMV-infected MEF by treating cells with thapsigargin (Tg), a potent activator of the UPR. Consistent with previously published data (21), IRE1-mediated *Xbp1* splicing was massively reduced in MCMV-infected WT MEFs at late times post infection (Fig. 5C). In contrast, *Xbpl*splicing was only minimally affected by MCMV in *Sel1L* ko MEF. These results demonstrated that MCMV cannot downregulate IRE1 protein levels and IRE1 signaling in the absence of SEL1L.

**Figure 5.**
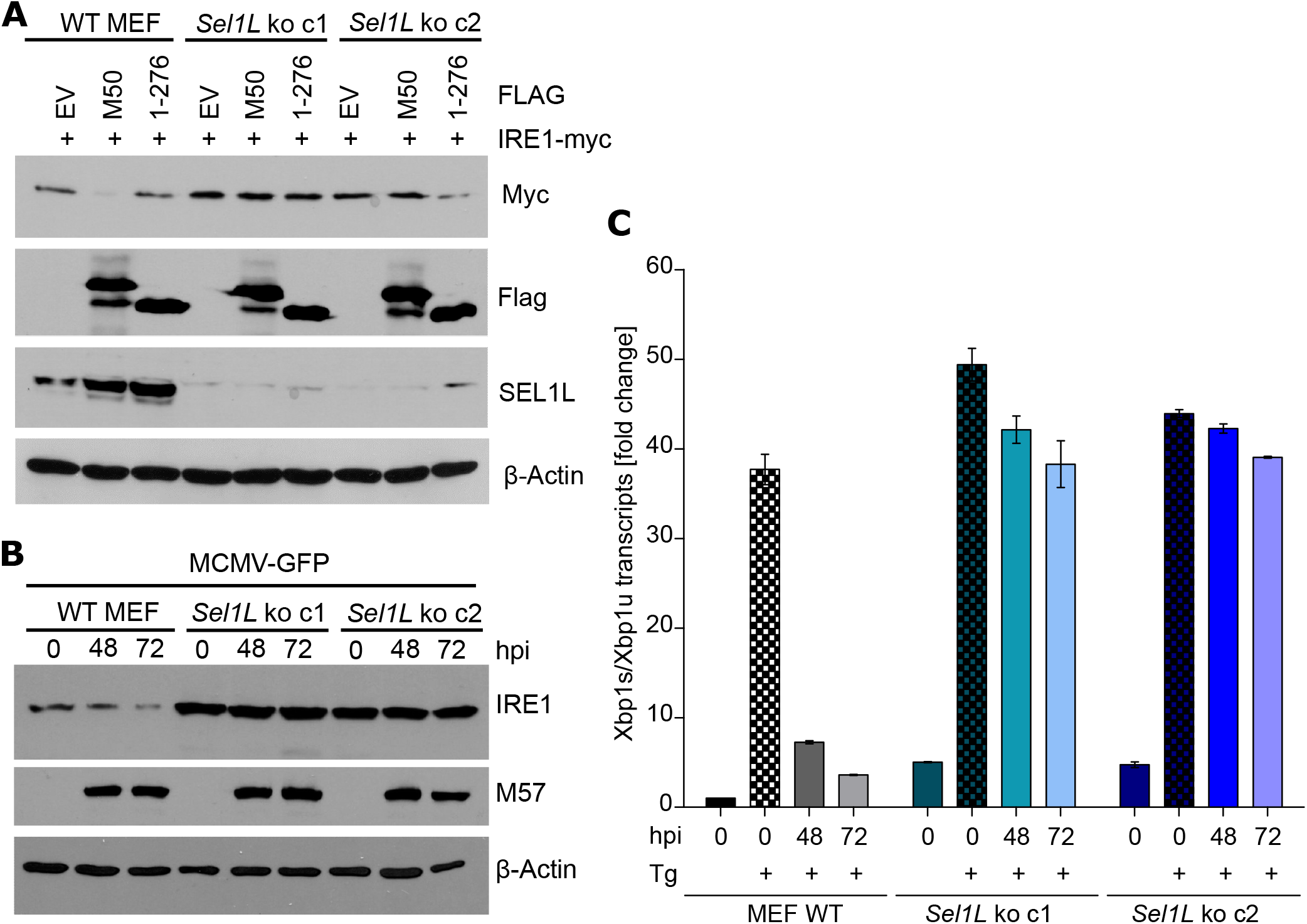
M50-mediated IRE1 degradation is abolished in SEL1L-deficient cells. (A) WT and *Sel1L* knockout MEFs were transfected with plasmids expressing myc-tagged IRE1 and full-length or truncated (1-276) M50 or empty vector (EV). Cell lysates were harvested 48 hours post transfection. (B) WT and *Sel1L* knockout MEFs were infected with MCMV-GFP (MOI=3). Cell lysates were analyzed by immunoblot. M57 was detected as an infection control. (C) WT and *Sel1L* knockout MEFs were MCMV-infected as in B and treated for 5 h with 60 nM thapsigargin (Tg). Total RNA was isolated and mRNA levels of *Xbpls* and *Xbplu* were determined by qRT-PCR. *Gapdh* was used for normalization. Fold changes relative to untreated WT MEFs are shown as means ±SEM of three biological replicates.

### *Sel1L* knockout affects viral gene expression

We have recently shown that MCMV briefly activated IRE1-mediated signaling at early times post infection to boost the activation of the viral major immediate-early promoter (20). At late times post infection, MCMV M50 downregulates IRE1 (21), presumably to avoid effects of the UPR that are detrimental for the virus. As SEL1L deficiency results in increased IRE1 levels, increased *Xbp1* splicing, and an inability of MCMV to downregulate IRE1 and inhibit IRE1 signaling at late times post infection (Fig. 5), we wanted to determine how SEL1L deficiency affects MCMV gene expression at early (2-8 hpi) and late (24-72 hpi) times post infection. As shown in Fig. 6, expression of the viral immediate-early 1 (IE1) protein was increased in *Sel1L* ko MEF at early times, but expression of viral early (M57) and late (glycoprotein B) proteins at later times was reduced or delayed. These results suggested that SEL1L deficiency impairs viral protein expression at late times post infection.

**Figure 6.**
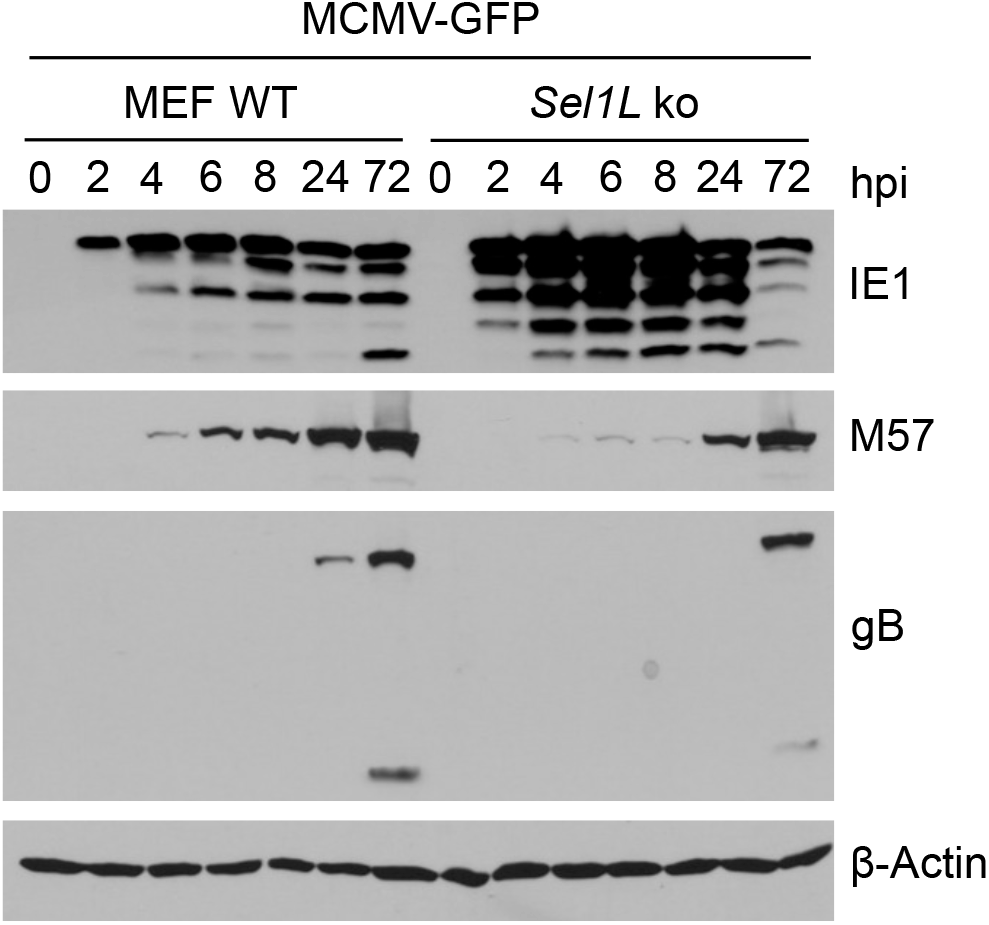
*Sel1L* knockout affects viral gene expression. WT and *Sel1L* knockout MEFs were infected with MCMV-GFP (MOI=3). Cell lysates were harvested at different times post infection. Expression of the immediate-early 1 (IE1), the early M57 protein, and the late glycoprotein B (gB) were analyzed by immunoblot.

## Discussion

In this study, we show that M50 interacts with IRE1 and SEL1L and induces IRE1 degradation via the canonical SEL1L-HRD1 ERAD pathway. Thus, M50 functions as a viral adaptor protein that tethers IRE1 to the SEL1L-HRD1 complex, thereby accelerating its decay (Fig. 7). The crucial role of SEL1L in M50-mediated IRE1 degradation was demonstrated with *Sel1L* ko cells, in which M50-mediated IRE1 degradation was abolished (Fig. 5). In these cells, viral protein expression was impaired at late times post infection (Fig. 6). This observation suggests that M50-mediated IRE1 degradation is beneficial for MCMV to secure viral protein expression in the late phase of infection. However, one has to consider that IRE1 is only one of many endogenous substrates of the SEL1L-HRD1 ERAD pathway. Thus, *Sel1L* knockout will most certainly also affect the turnover of many other cellular proteins. It has been shown that an acutely induced *Sel1L* knockout leads to an ERAD defect and a dilated ER (23). Studies with permanently SEL1L-deficient cells must therefore be interpreted with caution as these cells likely activate compensatory pathways to restore ER homeostasis and preserve cell viability. Proteasome inhibitors and inhibitors of the AAA ATPase p97 are even less useful for studying the biological importance of M50-induced IRE1 degradation as they affect many cellular functions besides ERAD. The best way to address this important question would be to use an M50 mutant that has selectively lost the ability to interact with SEL1L or with IRE1, but is otherwise fully functional, particularly with respect to M50’s function in nuclear egress of capsids. Unfortunately, such an M50 mutant has not been identified yet.

**Figure 7.**
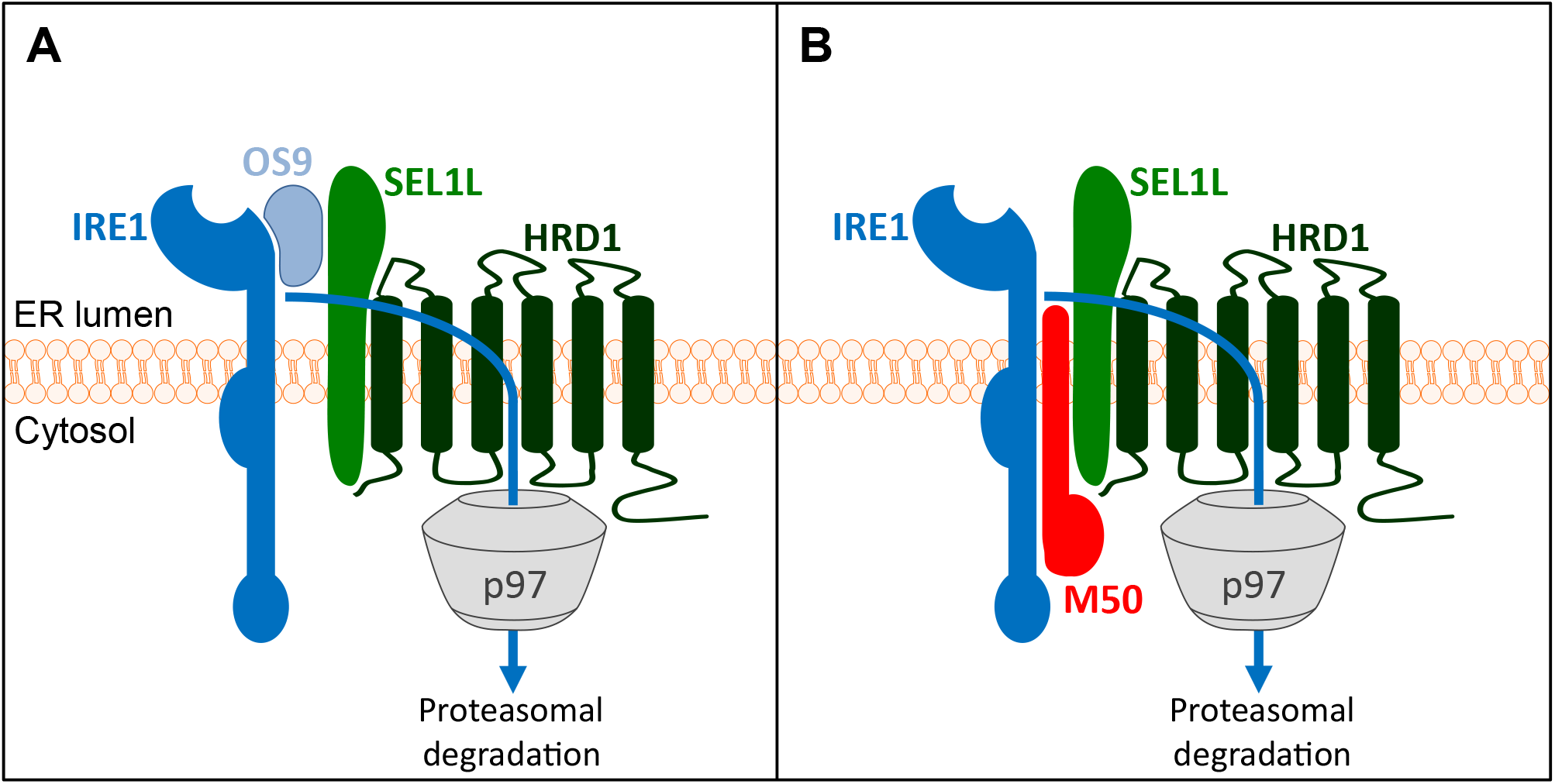
M50-mediated IRE1 degradation via ERAD (model) (A) The natural turnover of IRE1 involves binding to the cellular substrate recognition factor OS9, which recruits misfolded proteins to the SEL1L-HRD1 complex. IRE1 is ubiquitylated by HRD1 and dislocated into the cytosol by the AAA ATPase p97/VCP. (B) The MCMV M50 protein functions as a viral adaptor protein that tethers IRE1 to SEL1L, thereby promoting IRE1 degradation.

The SEL1L-HRD1 complex is the most conserved and best-characterized ERAD complex in mammalian cells. However, several other ERAD complexes have been described, which use different E3 ubiquitin ligases such as GP78, MARCHF6, RNF5, RNF103, RNF170, TMEM129, TRC8, TRIM13, or ZNRF4 (3, 8). A number of viruses exploit the ERAD pathway for the purpose of immune evasion or to provide an environment conducive to viral replication (24). The HCMV US2 and US11 proteins have been instrumental in studies of the mammalian ERAD system. They exploit separate components of the ERAD system to degrade MHC class I heavy chains, thereby impeding recognition of infected cells by cytotoxic T lymphocytes (25). While US2 appropriates the E3 ubiquitin ligase TCR8 to ubiquitylate MHC-I heavy chains (26), US11 uses a complex consisting of Derlin-1 and the E3 ligase TMEM129 (27, 28). Earlier studies have reported an association of US11 with SEL1L and concluded that US11 might induce MHC-I degradation via the canonical SEL1L-HRD1 complex (29). However, more recent work demonstrated that US11 promotes MHC-I degradation through the TMEM129- Derlin-1 complex, whereas US11 itself is subject to degradation via the SEL1L-HRD1 complex (27, 28).

M50 and UL50 are type II transmembrane proteins with a C-terminal membrane-spanning domain. In virus-infected cells, these proteins localize to the ER and the nuclear envelope, which is continuous with the ER. M50 and UL50 associate with the nuclear proteins M53 and UL53, respectively, leading to their accumulation at the inner nuclear membrane, were they recruit viral or cellular kinases to disrupt the nuclear lamina and facilitate nuclear egress of viral capsids. This important function of M50/UL50 and their respective partner proteins is highly conserved among the *Herpesviridae* and has been investigated extensively (reviewed in (30)). Apart from their function in nuclear egress, M50 and UL50 have additional functions such as the ability to interact with the UPR sensor IRE1 and induce its degradation (21). We show here that the MCMV M50 protein triggers degradation of IRE1 via the canonical SEL1L-HRD1 ERAD pathway. It is the same pathway, which the cell uses for IRE1 degradation during the natural turnover of IRE1 (9). This function requires interaction with SEL1L, a component of the canonical SEL1L-HRD1 ERAD complex (Fig. 5).

The HCMV UL50 protein can also induce IRE1 degradation in a similar way as M50 does (21). It seems likely that this occurs by the same or a very similar mechanism as the one we describe here for M50. However, this remains to be formally proven. Interestingly, two recent studies have shown that UL50 interacts with the ERAD machinery. It interacts with the ubiquitin-like modifier-activating enzyme 7 (UBA7, also known as UBE1L) and the ER-associated ubiquitin ligase RNF170 to induce ubiquitylation and proteasomal degradation of UBE1L (31). RNF170 is an as-yet poorly characterized E3 ubiquitin ligase involved in ERAD of the inositol trisphosphate receptor (32). It is unknown whether RNF170 can use SEL1L as a cofactor as HRD1 does. Nevertheless, it remains possible that RNF170 is somehow involved in the degradation of IRE1. A recent study has shown that UL50 also interacts with p97/VCP and downregulates p97 protein levels (33). How exactly UL50 downregulates p97 remains unclear. Interestingly, a short isoform of UL50 lacking the N-terminal 198 amino acids counteracts the downregulation of p97 by the full-length UL50 protein (33). This autoregulatory mechanism appears to be important for viral gene expression, probably because p97/VCP affects the splicing of the major immediate-early transcripts and the expression of the major viral transactivator protein, IE2 (34).

UL148 is another HCMV protein that directly affects UPR signaling. It triggers activation of PERK and IRE1 and remodels the ER (35, 36). Moreover, it regulates the composition of the viral gH-gL glycoprotein complex by interacting with the immature gH-gL complex (37) and by increasing the stability of gO, which is rapidly degraded via ERAD. Stabilization of gO involves interaction of UL148 with SEL1L, suggesting that UL148 dampens the activity of the SEL1L-HRD1 ERAD complex (38). Thus, interaction of viral proteins with SEL1L can either dampen ERAD, as shown for UL148, or promote ERAD, as shown here for M50.

## Materials and Methods

### Cells and virus

The following cell lines were used: immortalized MEFs (39), 10.1 fibroblasts (40), HEK 293A (Invitrogen) and HEK 293T cells (ATCC, CL-11268). Cells were grown under standard conditions in Dulbecco’s modified Eagle’s medium supplemented with 10% fetal calf serum, 100 IU/ml penicillin and 100 μg/ml streptomycin (Sigma).

MCMV-GFP (41) was propagated and titrated on 10.1 fibroblasts. Viral titers were determined by using the median tissue culture infective dose (TCID50) method.

### Plasmids and transfection

Plasmids pcDNA3-mIRE1-3xmyc, hIRE1-HA, M50-Flag, and M50(1-276)-Flag were described previously (21). A pcDNA3.1 plasmid encoding HA-tagged K48-only Ub (22) was kindly provided by Vishva Dixit (Genentech). The murine *Sel1L* gene was PCR-amplified from pCMV6-Sel1L-Myc-DDK (OriGene) and subcloned in pcDNA3. HEK-293A cells were transiently transfected polyethylenimine (Sigma), and MEFs were transfected using GenJet (SignaGen).

### CRISPR/Cas9 gene editing

CRISPR/Cas9 gene editing was used to generate *Sel1L* knockout MEFs essentially as described (20, 42). Briefly, two different guide RNAs (5’-GTCGTTGCTGCTGCTCTGCG-3’ and 5’-GCTGCTCTGCGCGGTGCTCC-3’) targeting the murine *Sel1L* gene were designed using E-CRISP (http://www.e-crisp.org/E-CRISP/designcrispr.html) and inserted into the lentiviral vector pSicoR-CRISPR-puroR (kindly provided by R. J. Lebbink, University Medical Center Utrecht, Netherlands). Lentiviruses were produced in HEK-293T cells using standard third-generation packaging. Lentiviruses were used to transduce MEFs in the presence of 5 μg/ml polybrene (Sigma). Cells were selected with 1.5 μg/ml puromycin (Sigma), and single cell clones were obtained by limiting dilution.

### Immunoprecipitation and immunoblot analysis

Whole cell lysates were obtained by lysing cells in RIPA buffer supplemented with a cOmplete Mini protease inhibitor cocktail (Roche). Protein concentrations were measured using a BCA assay (Thermo Fisher Scientific). Equal protein amounts were boiled in sample buffer and subjected to SDS-PAGE and semi-dry blotting onto nitrocellulose membranes. For immunodetection, antibodies against the following proteins and epitopes were used: HA (16B12, BioLegend), β-actin (AC-74, Sigma), IRE1 (14C10, Cell Signaling), SEL1L (151545, Biomol). Monoclonal antibodies against MCMV IE1 (CROMA101), M57 (M57.02), and M55/gB (SN1.07) were from the Center for Proteomics, University of Rijeka, Croatia. Secondary antibodies coupled to horseradish peroxidase (HRP) were purchased from Dako. Detection of IRE1 ubiquitination was done as described in detail elsewhere (43).

### RNA isolation and quantitative PCR

Total RNA was isolated from WT and *Sel1L* ko MEFs using an innuPREP RNA Mini Kit (Analytik-Jena). Reverse transcription and cDNA synthesis was carried out with 2 μg RNA using 200 U RevertAid H Minus Reverse Transcriptase, 100 pmol Oligo(dT)18, and 20 U RNase inhibitor (Thermo Fisher Scientific). Quantitative real-time PCR reactions employing SYBR Green were run on a 7900HT Fast Real-Time PCR System (Applied Biosystems). PCR primers for *Xbpls, Xbplu, Gapdh*, and *Ire1* have been described (20, 21). *Chop* was amplified with primers 5’-TATCTCATCCCCAGGAAACG-3’ and 5’-GGGCACTGACCACTCTGTTT-3’. Reactions were performed under the following conditions: 45 cycles of 3 sec at 95°C and 30 sec at 60°C. Three replicates were analyzed for each condition, and the relative amounts of mRNAs were calculated from the comparative threshold cycle (Ct) values by using *Gapdh* as reference.

### Inhibitors

The proteasome inhibitor MG-132 was purchased from Sigma. The AAA ATPase Inhibitor CB-5083 was from Xcessbio.

### Statistical analysis

All statistical analyses were performed with GraphPad Prism 5.0 software. One-way ANOVA followed by Bonferroni’s post hoc test was used for the analysis of qPCR experiments.

## Acknowledgements

We thank Elena Muscolino and Olha Puhach for critical readings of the manuscript.

This study was supported by the Deutsche Forschungsgemeinschaft (grant BR1730/6-1 to W.B.). The Heinrich Pette Institute is supported by the Free and Hanseatic City of Hamburg and the Federal Ministry of Health. The funders had no role in study design, data collection and interpretation, or the decision to submit the work for publication.

